# Characterisation of membrane protein interactions by peptidisc-mediated mass photometry

**DOI:** 10.1101/2023.06.01.543083

**Authors:** John William Young, Emanuel Pfitzner, Philipp Kukura, Carol V. Robinson

## Abstract

Membrane proteins perform a variety of critical functions in the cell, making many of them primary drug targets. At the same time, their preference for a lipid environment makes them challenging to study using established solution-based methods. Here, we show that peptidiscs, a recently developed membrane mimetic, provide an ideal platform to study membrane proteins and their interactions with mass photometry (MP) in detergent-free conditions. The mass resolution for membrane protein complexes is similar to that achievable with soluble proteins owing to the low carrier heterogeneity. Using two well-characterized bacterial membrane protein complexes - the ABC transporter BtuCD, and the Sec translocon - we show that MP can quantify interactions between peptidisc-reconstituted membrane protein receptors and their soluble protein binding partners. Using the BAM complex, we further show that MP reveals interactions between a membrane protein receptor and a bactericidal antibody. Our results highlight the utility of peptidiscs for membrane protein characterization in detergent-free solution and provide a rapid and powerful platform for quantifying membrane protein interactions.

## INTRODUCTION

Integral membrane proteins are a highly relevant class of biological molecules, and control many of the essential processes of life including energy production, nutrient import, cell-cell signalling, protein translocation and comprise 60% of current drug targets ^1-4^. However, membrane proteins are notoriously difficult to study because of their relatively low abundance and high hydrophobicity. Before being analysed by most structural and/or biochemical techniques, membrane proteins must first be removed from hydrophobic cellular membranes ^5^. Membrane protein extraction from cellular membranes is most often achieved using detergents, which solubilize the lipid bilayer and maintain membrane proteins in a soluble state by shielding their hydrophobic surfaces from water, thereby preventing aggregation. However, even the mildest detergents can disrupt native protein conformations and/or dissociate transiently associated complexes, leading to loss of potentially relevant protein-protein and protein-lipid interactions ^6-10^.

To minimize these detergent effects, numerous membrane mimetic systems have been developed in recent years, which stabilize membrane proteins in a water-soluble format by shielding their hydrophobic transmembrane regions from the aqueous environment. Commonly used membrane mimetics for structural and/or biochemical analysis of membrane proteins include nanodiscs, saposins, peptidiscs, and SMALPs ^1, 11-13^. With different membrane mimetic systems available, how do researchers identify the optimal conditions for reconstituting their protein of interest in detergent-free conditions? This is often a non-trivial task and can involve applying different reconstitution strategies before identifying optimal conditions for downstream structural or biochemical analysis. Post-reconstitution sample quality is often screened using size exclusion chromatography (SEC), light scattering coupled to SEC (SEC-MALS), analytical ultracentrifugation (AUC), and negative stain electron microscopy (NS-EM) ^14-16^. Although successful, these techniques are time consuming, relatively low throughput and, apart from NS-EM, consume substantial amounts of valuable protein sample ^14^. Mass Photometry (MP), single molecule mass measurement in solution, has recently addressed many of these limitations for soluble proteins and has emerged as a powerful approach for characterising protein-protein interactions ^17-19^. When combined with membrane mimetics including nanodiscs and SMALPs, MP can be very useful to rapidly assess the homogeneity of reconstituted membrane protein samples ^20, 21^. However, the utility of MP for quantifying interactions between reconstituted membrane proteins and their soluble protein binding partners has not yet been explored. In addition, the achievable mass resolution in MP ultimately hinges on the homogeneity of the chosen carrier, which is substantial for most membrane mimetics.

Recently, peptidiscs have emerged as an alternative membrane protein carrier featuring a particular straightforward purification procedure ^9, 12, 22^. Here, we use a series of His-tagged bacterial integral membrane protein complexes to demonstrate that membrane protein complexes reconstituted in peptidiscs are amenable to analysis by MP, while exhibiting minimal heterogeneity despite a rapid and simple purification procedure (Figure 1). Using the ABC transporter BtuCD and the SecYEG protein-conducting channel as examples, we demonstrate that MP can be used to quantify high-affinity binding interactions between peptidisc-reconstituted membrane protein receptors and their naturally occurring soluble protein ligands. We further use MP to characterize binding of a bactericidal antibody onto the outer membrane-embedded pentameric BAM complex. These results highlight the utility of the combination of MP with peptidiscs for quantifying protein-protein interactions at the single molecule level.

**Figure 1:**
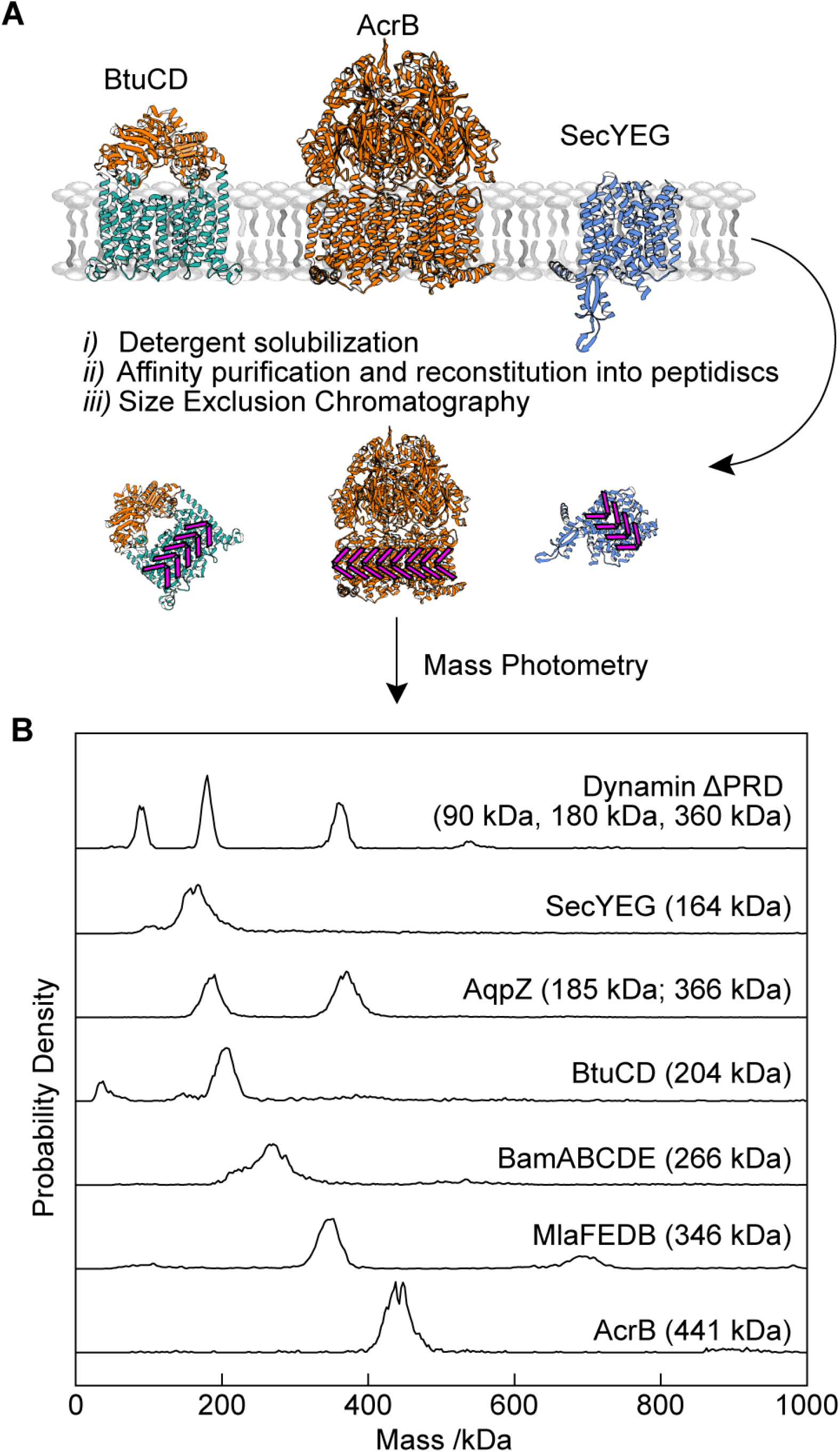
Overview of experimental workflow. **(A)** The overexpressed target protein is extracted from biological membranes with mild detergent (DDM) prior to affinity purification and reconstitution into peptidiscs. Immediately following SEC purification, peptidisc particles are analysed by Mass Photometry (MP). **(B)** Representative mass histograms for each peptidisc-reconstituted integral membrane protein analysed in this study. The top panel (labelled “Dynamin ΔPRD”) represents the soluble protein standard used to calibrate the MP instrument during these measurements.

## MATERIALS AND METHODS

### Plasmid preparation and protein expression

Plasmids for expression of *E. coli* BtuCD, BtuF, AcrB and BamABCDE were obtained from our laboratory collection ^23, 24^. BtuF and AcrB are 6x His-tagged on their C-termini; BtuCD is 10x His-tagged on the BtuC subunit. To facilitate expression of *E. coli* AqpZ, our laboratory’s existing pET15b-AqpZ-GFP construct was modified using the Polymerase Incomplete Primer Extension (PIPE) method to delete the GFP sequence and add a C-terminal 6x His-tag onto AqpZ ^25, 26^. The plasmid for expression of *E. coli* SecYEG has been previous described ^27^. The plasmid for expression of E. coli MlaFEDB was a kind gift from Dr. Damien Ekiert (NYU) ^28, 29^. The Syd open reading frame (containing amino acids 1-181) was PCR-amplified from *E. coli* BL21 (DE3) genomic DNA and inserted into the pET-22b expression vector (Novagen) between the NcoI and XhoI cloning sites using an In-Fusion cloning kit (Clonetech). All plasmids were amplified by transforming them into *E. coli* Stellar Competent Cells (Takara) and the DNA sequences were verified by Sanger sequencing.

For each protein, the appropriate plasmid was transformed into chemically competent *E. coli* C43(DE3) cells (Cambridge Bioscience). A single freshly transformed colony was inoculated into 40 ml LB media supplemented with 100 mg/mL Ampicillin and grown overnight at 37 °C. The next morning, the preculture was diluted 1/100 into 4 L fresh LB media (plus antibiotic) and grown at 37 °C until the culture reached OD600 nm (OD600) between 0.4 and 0.6. Protein production was induced by addition of either 0.2% Arabinose (for SecYEG and MlaFEDB) or with 0.5 mM Isopropyl-b-D-1-thiogalactopyranoside (IPTG – for all other constructs), and cultures were grown a further 3 hours at 37□°C. Cells were collected by centrifugation at 5,000xg for 10 min at 4□°C in a Beckman JLA 8.1000 rotor. Excess media was discarded, and cell pellets were resuspended in 20 mL Buffer A (20 mM Tris HCl pH 8, 150 mM NaCl) and stored at -80 °C until use. The antibody mAB1 (anti-BamA monoclonal antibody 15c9) was obtained as a lyophilized powder from Genentech under Material Transfer Agreement OR-216904. The powder was resuspended in PBS buffer to a concentration of 5 mg/mL and stored at -80 °C until use.

### Purification of soluble proteins BtuF and Syd

Resuspended cells were homogenized by gentle stirring and supplemented with an EDTA-free protease inhibitor cocktail (Roche). The cell suspension was passed three times through an M-110 PS microfluidizer (Microfluidics) at 15,000 psi. Following cell lysis, insoluble material was pelleted by centrifugation at 20,000xg for 20 min at 4□°C in a JA 25.50 rotor. The supernatant was loaded onto a home-packed 5 mL Ni-NTA column equilibrated in Buffer B (Buffer A supplemented with 25 mM Imidazole) and allowed to pass via gravity flow. The column was washed first with 50 mL of Buffer B, then with 25 mL of Buffer C (Buffer A supplemented with 80 mM Imidazole). Bound proteins were eluted in 20 mL Buffer A supplemented with 250 mM Imidazole. After verifying sample purity by SDS-PAGE, fractions containing the target proteins were pooled, concentrated, and further purified by size exclusion chromatography (SEC) on a Superdex 200 10/300 column equilibrated in Buffer A. Peak fractions were pooled and stored at -80 °C until use.

### Purification of membrane proteins and reconstitution into peptidiscs

Resuspended cells were homogenized by gentle stirring and supplemented with an EDTA-free protease inhibitor cocktail (Roche). The cell suspension was passed three times through an M-110 PS microfluidizer (Microfluidics) at 15,000 psi. Following cell lysis, insoluble material was pelleted by centrifugation at 20,000xg for 20 min at 4□°C in a JA 25.50 rotor. To pellet cellular membranes, the supernatant was ultracentrifuged at 200,000xg for 30 minutes at 4□°C in a Beckman SW32Ti rotor. Membrane protein purification and reconstitution into peptidiscs was performed as previously described with minor modifications ^22,9^. Membranes were resuspended in 20 mL Buffer A and solubilized with 1% n-Dodecyl-β-D-Maltopyranoside (DDM) for 30 minutes at 4□°C. Insoluble material and aggregates were removed by centrifugation (20,000xg, 20 minutes). The detergent-solubilized material was then loaded onto a home-packed 5 mL Ni-NTA column equilibrated in Buffer A supplemented with 0.02% DDM and allowed to pass via gravity flow. The column was washed first with 50 mL of Buffer B supplemented with 0.02% DDM, then with 25 mL of Buffer C supplemented with 0.02% DDM. To facilitate reconstitution into peptidiscs, the resin was resuspended in 50 mL of Buffer A supplemented with 1 mg/mL peptidisc peptide. Peptidisc peptides were obtained as a lyophilized powder from Peptidisc Biotech ^12^. After collecting excess peptides, the resin bed was washed again with 50 mL of Buffer A. Reconstituted proteins were then eluted in Buffer A supplemented with 250 mM Imidazole. After analysis by SDS-PAGE to verify sample purity, peak fractions were pooled, concentrated, and injected onto a Superdex 200 GL 10/300 column equilibrated in Buffer A. Peak fractions were collected. Prior to MP analysis, protein concentration was determined using UV-vis spectroscopy by monitoring the absorbance at 280 nm. To determine the theoretical extinction coefficients for each membrane protein used in this study, the protein amino acid sequence along with the sequence of the peptidisc peptide scaffold was inputted into the Expasy ProtParam tool. The resulting extinction coefficient values are listed in Table 1 below.

**Table 1:** Molar extinction coefficients for the peptidisc-reconstituted membrane proteins used in this study.

| **Membrane Protein** | **Extinction Coefficient ($M^{-1}cm^{-1}$)**<br>**in peptidiscs** |
| --- | --- |
| AcrB | 103710 |
| MlaFEDB | 92820 |
| BtuCD | 120430 |
| AqpZ | 48930 |
| SecYEG | 84800 |
| BamABCDE | 310100 |

### Mass Photometry analysis of reconstituted membrane proteins

Mass Photometry measurements were performed on cleaned glass cover slips and recorded on a mass photometer (TwoMP, Refeyn Ltd.) as previously described ^17, 20^. In a typical experiment, we first added 5 μL of clean PBS buffer to a silicone gasket (Grace Bio-Labs reusable CultureWell™ gaskets, 50 wells, 3 mm x1mm, Merck Life Science UK Limited) mounted on the clean coverslip to find the glass/buffer interface. Following SEC purification, successive fractions of each reconstituted membrane protein were diluted 100-fold in clean PBS buffer to a concentration of ∼5-20 nM. 20 μL of the dilution were added to the 5 μL PBS buffer and a 60 s movie was immediately recorded. The movies were analysed using DiscoverMP software (version 2.5) to quantify protein binding events. Sample molecular weights were obtained by contrast comparison with known mass calibrants measured on the sample day (Figure S6). The resulting events were then further analysed and plotted with home written Python scripts using the libraries NumPy, matplotlib, and SciPy ^30-32^. All mass spectra are kernel density estimates of the resulting histograms with the kernel being a Gaussian of width σ = 2.5 kDa. Where necessary, distributions were fitted with a sum of Gaussians:

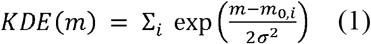

To quantify interactions between BtuCD and BtuF, a series of 10 binding mixtures was prepared in 10 μL volume using PBS plus 5 mM MgCl_2_ as the buffer. The concentration of BtuCD was held constant at ∼200 nM across all mixtures, and the concentration of BtuF was increased from 0 – 1200 nM. Following incubation at room temperature for 30 minutes, each mixture was analysed by MP. Immediately before measuring, 2 μL of each was diluted into 20 μL clean PBS and analysed exactly as described above. To test the effect of ATP on the BtuCD-F interaction, the binding titration was repeated with 1 mM ATP present in all conditions. Measurements were repeated multiple times on different days to ensure reproducibility. A similar protocol was used to quantify interactions between SecYEG and Syd, with two modifications: 5 mM MgCl_2_ was omitted from the buffer, and the concentration of Syd was varied between 0-500 nM.

The fitted masses or centre of masses of the titration datasets shown in figures 4 and 5 were fitted with a Hill equation to extract an estimate for the dissociation constant:

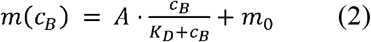

To monitor interactions between the BAM complex and mAB1, a series of 8 binding mixtures was prepared in 10 μL volume, using PBS as the buffer. The concentration of BAM was fixed at 400 nM, and the concentration of mAB1 was increased from 0-600 nM. Binding mixtures were analysed as described above.

## RESULTS AND DISCUSSION

### Peptidisc reconstitution of bacterial membrane proteins and analysis by Mass Photometry

To establish our method, we started with the *E. coli* 340 kDa trimeric drug efflux pump AcrB ^**33**^. After over-expressing His-tagged AcrB and solubilizing the membrane fraction with DDM, we purified the protein and reconstituted it into peptidiscs (Materials and Methods). Following SDS-PAGE analysis of the purified material (Figure 2A), fractions were pooled, concentrated, and further purified by size-exclusion chromatography (SEC) in detergent-free buffer (Figure 2B). Several fractions under the main peak (labelled Fractions 1-4) were collected and each was analysed by MP. Particle landing events were recorded as movies and analysed using commercial software (DiscoverMP, Refeyn Ltd) to produce mass histograms (Figure 2C). We found little variation between individual fractions, with each containing a major species centred at ∼ 430-440 kDa. To ensure our method is consistent and reproducible, we performed this workflow twice, with two independently prepared batches of cells. Overlayed histograms from two independent experiments are shown (Figure 2D). The two replicates are reassuringly similar, with only minimal differences observed in the measured molecular weight. We note that previous studies on *E. coli* AcrB have shown that the protein co-purifies with a large number of annular lipids ^**34**, **35**^. Thus, it is likely that the 90-100 kDa difference in mass between our peptidisc preparation and the “naked” protein in detergent micelles arises from both peptidisc peptides and phospholipids.

**Figure 2:**
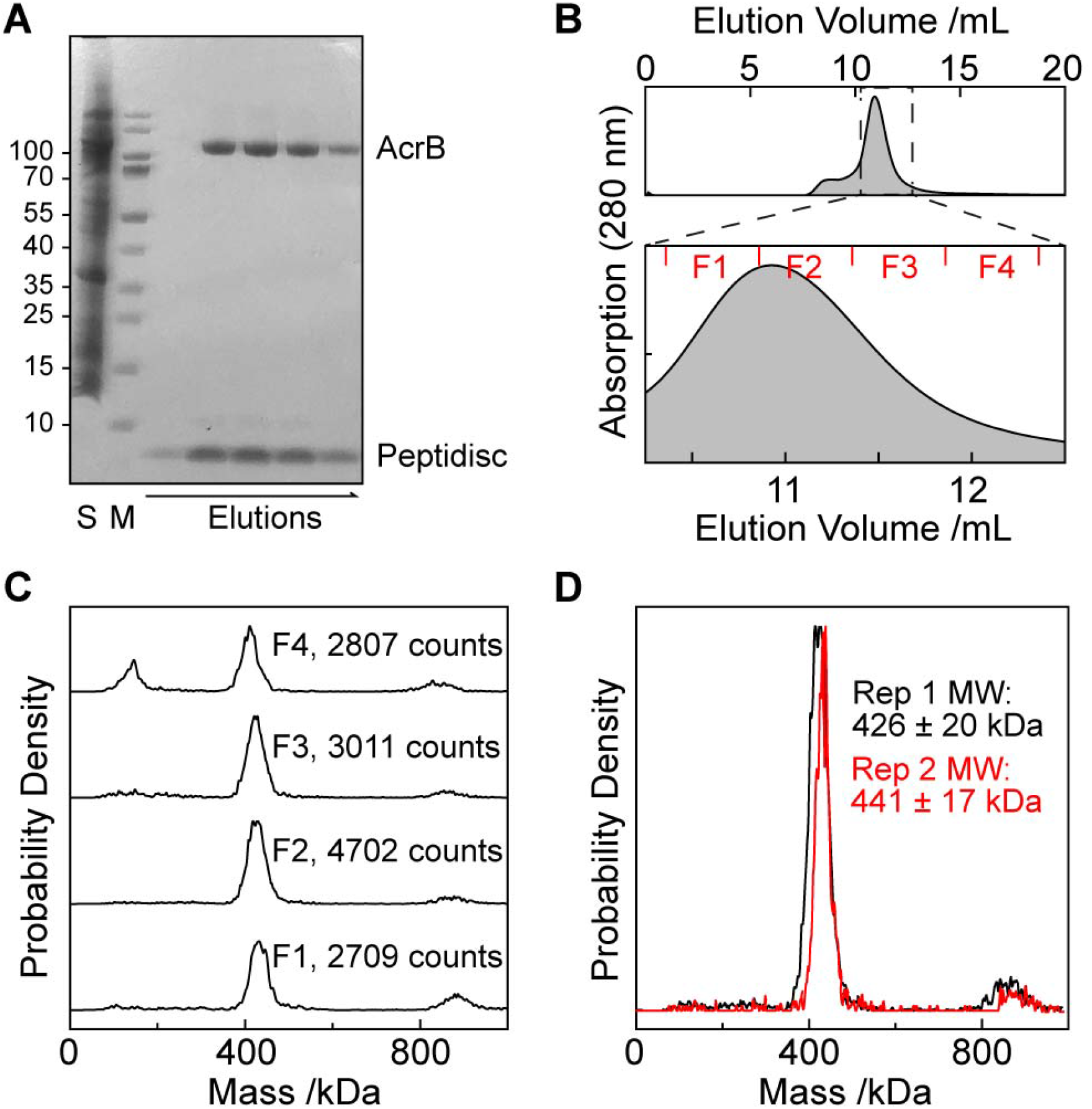
Mass Photometry of the trimeric drug efflux pump AcrB reconstituted in peptidiscs. **(A)** SDS-PAGE analysis of purified AcrB following affinity purification and peptidisc reconstitution. The un-purified detergent-solubilized membrane extract is shown in the left-most lane (labelled “S”); purified eluted protein is indicated by “Elutions”; prestained molecular weight marker is in lane “M”. **(B)** Size exclusion chromatography (SEC) trace and fractions from (A) on a Superdex 200 10/300 column. Protein absorbance as a function of elution volume was monitored at 280 nm. Fractions under the main peak (labelled F1-4) were collected and used in further analysis. **(C)** Each fraction from **B** analysed by MP. The total number of particles (or “landing events”) observed in each movie is indicated. **(D)** Representative fractions from two independent biological replicates.

Next, we applied the same workflow to the 260 kDa phospholipid ABC transporter MlaFEDB (Figure 3A-C) and the 130 kDa Vitamin B_12_ ABC transporter BtuCD (Figure 3D-F). Following SEC purification of the reconstituted protein complexes, successive fractions were analysed by MP. We observed only minimal differences between individual fractions for each complex. For MlaFEDB, each fraction contains a major species centred at ∼340 kDa (Figure 3B); for BtuCD, each fraction contains a major species centred at ∼215 kDa (Figure 3E) The quality of these reconstitutions is highly reproducible, with only minimal differences observed between biological replicates (Figure 3C and F). Previous studies on MlaFEDB and BtuCD have indicated that these complexes co-purify with phospholipids ^**23**, **29**^; thus, the ∼80 kDa difference in mass between our peptidisc preparations and the theoretical molecular weights of MlaFEDB and BtuCD can be attributed to both peptidisc peptides and phospholipids.

**Figure 3:**
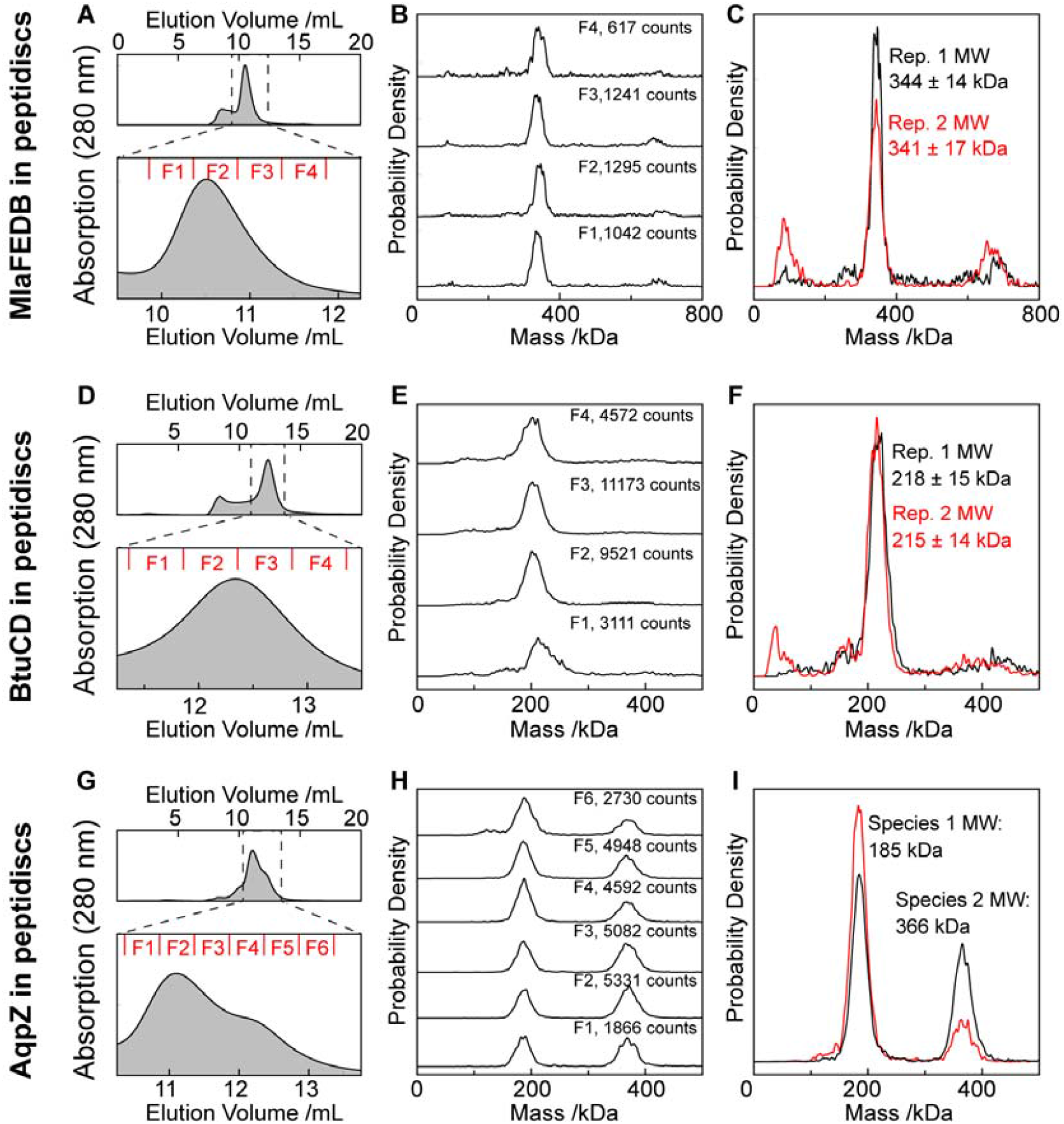
Mass Photometry of a selection of integral membrane proteins reconstituted in peptidiscs. **(A)** The ABC transporter MlaFEDB was purified and reconstituted into peptidiscs as in Figure 2. **(B)** MP analysis of the indicated fractions from **A**, including total number of detected particles per experiment. **(C)** Reproducibility from two independent biologicial replicates. **(D-F)** As in **A-C**, but for the ABC transporter BtuCD. **(G and H)** As above, but for the aquaporin AqpZ. **(I)** Fraction 5 from (H) was analyzed by MP in the absence (black trace) and presence (red trace) of 25 mM Imidazole.

When applying this same workflow to His-tagged *E. coli* aquaporin AqpZ (Figure 3G-I), we noticed that the SEC trace was somewhat broader than for the proteins described above (Figure 3G). When we analysed 6 successive fractions under the main peak we observed two distinct populations: a first sub-population centred at ∼180 kDa (54% abundance) and a second centred at ∼360 kDa (46 % abundance) (Figure 3H). Since the molecular weight of the second species is exactly double that of the first, we rationalized that this may represent two discs stacking together. Indeed, a recent report showed using a combination of SEC and electron microscopy that His-tags can induce formation of non-physiological “stacked” oligomers of membrane proteins ^36^. To test this possibility, we incubated one of the fractions with 25 mM Imidazole and repeated the measurement (Figure 3I). In the presence of Imidazole, we observed considerably less of the ∼360 kDa species (25% abundance), and a correspondingly higher amount of the 180 kDa species (75% abundance), confirming our hypothesis of His-tag induced dimerization occurring with AqpZ.

### Quantifying the interaction of BtuF and BtuCD using Mass Photometry

We next tested whether we could observe and quantify interactions between peptidisc-reconstituted receptors and their native protein ligands. We started with BtuCD, which interacts with the periplasmic B_12_ binding protein BtuF. Interactions between BtuCD and BtuF have been extensively characterized using detergent micelles and proteoliposomes ^37-39^. The structure of the BtuCD-F complex has been determined using X-ray crystallography, and the kinetics of the BtuCD-F interaction have been measured using Microscale Thermophoresis (MST) and Surface Plasmon Resonance (SPR) ^37-39^. These previous studies have shown that BtuF binds BtuCD with high affinity, and that the dissociation constant (K_D_) for this interaction is in the nanomolar range ^38, 39^.

Characterisation of BtuCD by MP revealed a species at ∼208 kDa (Figure 4A, black trace). The 5 kDa difference from our earlier measurement is well within the expected 2% RMS error for independent MP measurements ^18^. The measured mass increases to ∼230 kDa after incubation with a molar excess of BtuF, indicating that BtuF is binding to peptidisc-reconstituted BtuCD (Figure 4A, blue trace). Given that BtuF is a 30 kDa protein, a partial shift to 228 kDa indicates that not all available BtuCD complexes are bound to BtuF under these conditions. To increase the fraction of BtuCD bound to BtuF, we repeated the measurement in the presence of 1 mM ATP, which promotes BtuCD-BtuF interactions^39^, increasing the mass to 238 kDa.

**Figure 4:**
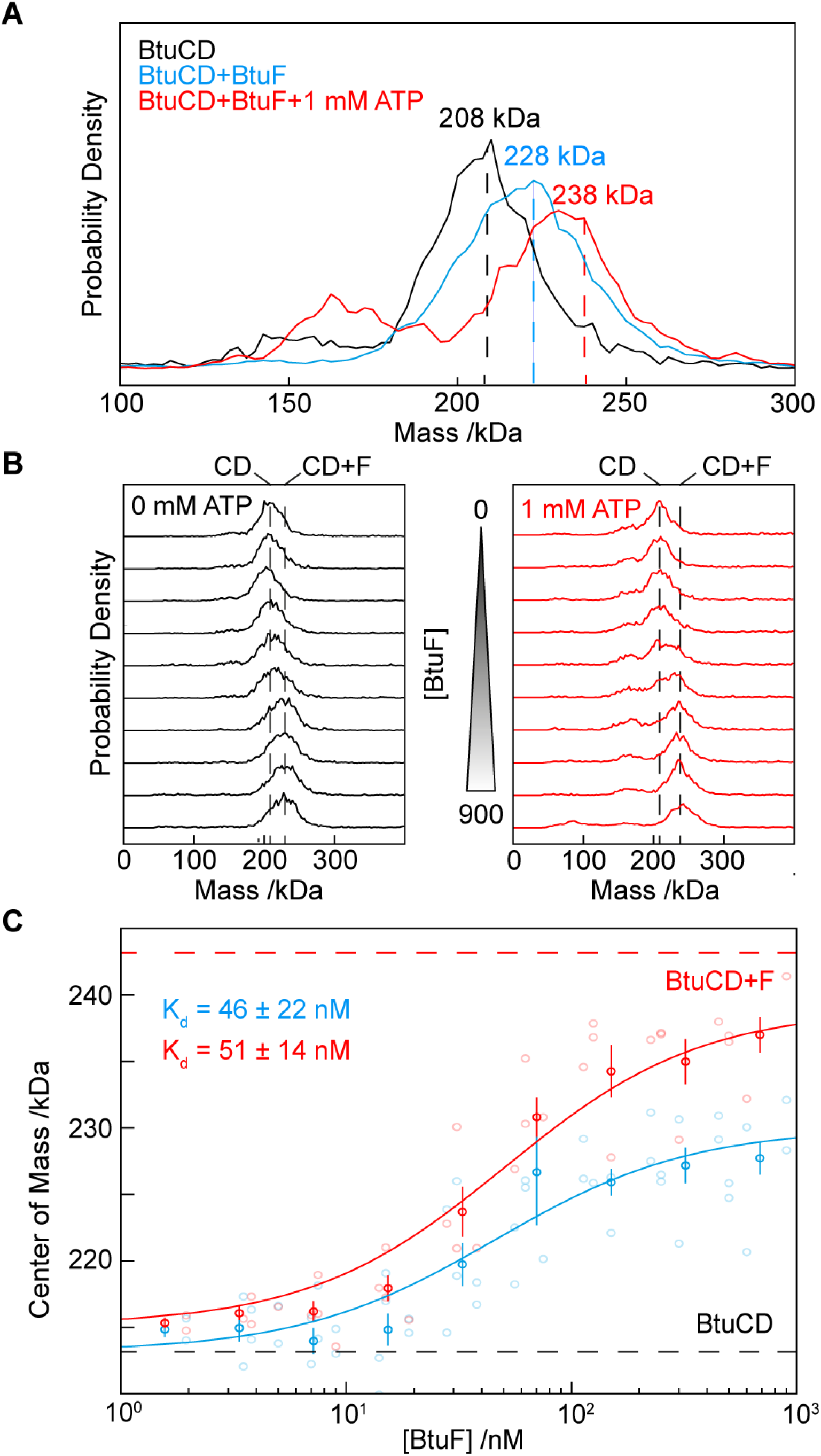
Quantifying interactions between the ABC transporter BtuCD and its protein binder BtuF. **(A)** A representative fraction under the main peak for BtuCD in the absence (black trace) or presence (blue trace) of a molar excess of BtuF. The “+BtuF” measurement was repeated in the presence of 1 mM ATP (red trace). The masses measured for each sample are indicated on the plot. **(B)** Incubations of aliquots of BtuCD with increasing amounts of BtuF in the absence (left panel) or presence (right panel) of 1 mM ATP. **(C)** The change in peak centre mass for each condition in **B** as a function of BtuF concentration. The data plot is derived from 6 independent experiments where the gray data points indicate the individual data and the black the averaged center of mass. The solid lines indicate Hill equation fitted to each dataset to derive the apparent K_D_ values for these conditions.

To quantify the binding of BtuF onto BtuCD in the presence and absence of ATP, we titrated BtuF from 0-1200 nM in the presence of BtuCD (Figure 4B). After an incubation for 30 minutes, each mixture was analysed by MP (Materials and Methods). We then plotted the center of mass of the resulting peaks (Figure 4B) as a function of BtuF concentration (Figure 4C), repeating the titrations repeated multiple times on different days to ensure reproducibility. Averaging measurements and fitting with a monovalent Hill equation (Equation 2) allowed us to derive the K_D_ for this interaction in the presence and absence of ATP. BtuF displays a similar affinity for BtuCD in the presence and absence of ATP, with both measured K_D_ values in the nanomolar range (Figure 4C). We find that ATP does not significantly increase the affinity of BtuF for BtuCD, but rather increases the fraction of BtuCD bound to BtuF.^39^

### Quantifying the interaction of Syd and SecYEG using Mass Photometry

Turning to a second example for which interaction kinetics and dissociation constant is not known we selected the bacterial SecYEG complex and its interaction with the 20 kDa cytosolic protein Syd ^40, 41^. Earlier experiments on the SecYEG-Syd interaction using SEC-MALS and non-denaturing gel electrophoresis suggest Syd binds to SecYEG with high affinity. We thus reconstituted SecYEG into peptidiscs and analysed the purified material by MP in the presence and absence of a molar excess of Syd. The measured mass for SecYEG alone is 164 kDa (Figure 5A, black trace). In the presence of Syd, the observed mass increases to 183 kDa, confirming that SecYEG and Syd interact strongly under our experimental conditions (Figure 5A, red trace). We next prepared a titration to quantify the SecYEG-Syd interaction. We fixed the concentration of SecYEG and increased the concentration of Syd from 0 - 500 nM (Figure 5B). After a brief incubation for 30 min, we analysed each solution by MP, determining the peak centre mass by fitting a Gaussian distribution to the histograms (Figure 5B) and plotting the result as a function of Syd concentration (Figure 5C). The resulting binding curve could be well-described by a monovalent Hill function (Equation 2), yielding a K_D_ = 194 ± 18 nM (Figure 5C).

**Figure 5:**
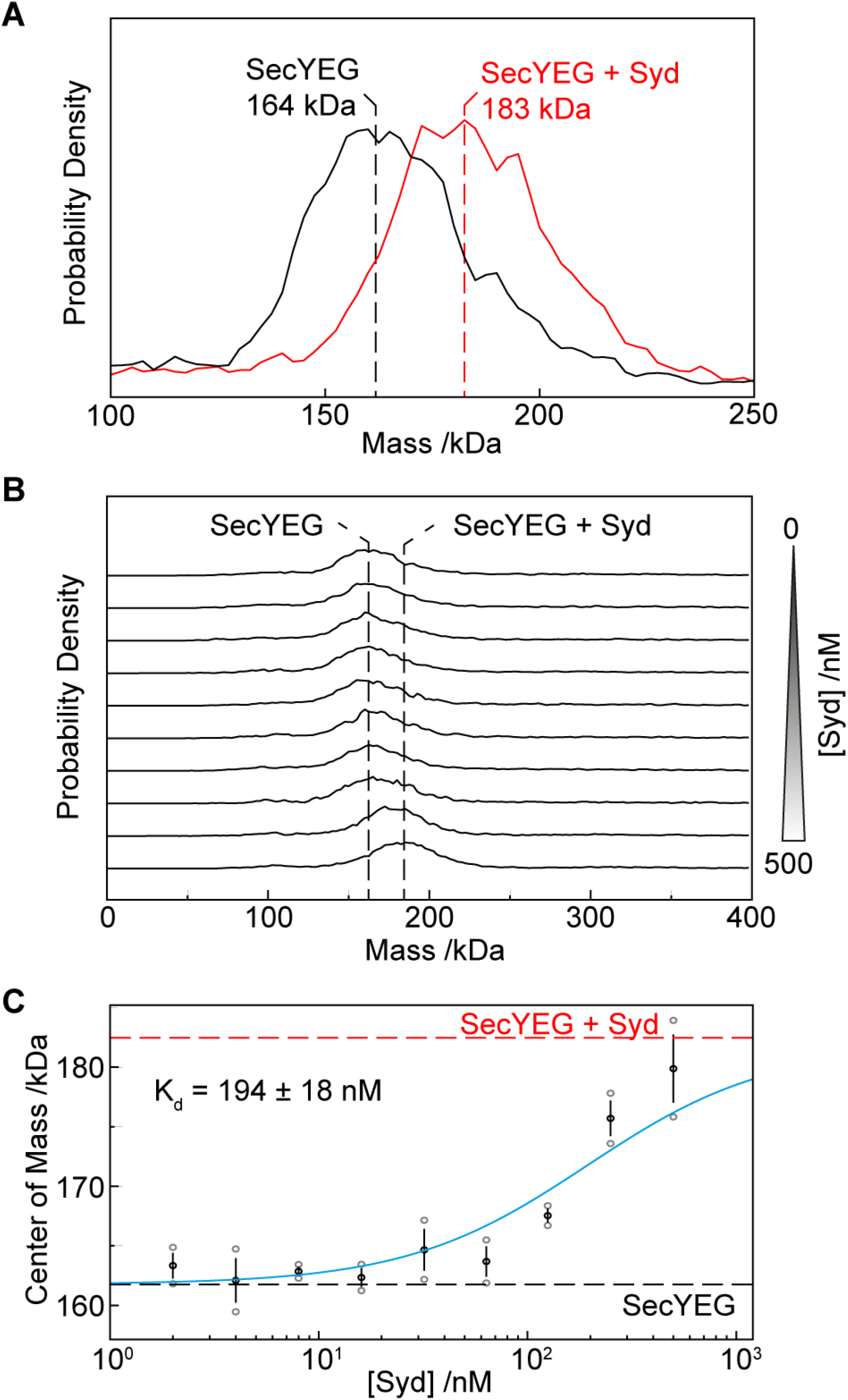
Quantifying interactions between the bacterial Sec translocon SecYEG and its protein binder Syd. **(A)** A representative fraction under the main peak for SecYEG in the absence (black trace) or presence (red trace) of a molar excess of Syd. The masses measured for each sample are indicated on the plot. **(B)** Aliquots of SecYEG incubated with increasing amounts of Syd. **(C)** The change in peak centre mass for each measurement in **B** plotted as a function of Syd concentration to yield the dissociation constant (K_D_). The blue line represents a Hill function fitted to the resulting masses to determine the dissociation constant for this interaction. The data points represent an average of derived from two independent experiments.

### Monitoring interactions between the BAM complex and a bactericidal antibody

Having verified that our approach is effective for monitoring high-affinity interactions between membrane proteins and their naturally occurring protein ligands, we next assessed whether our method can be extended to monitor interactions between a membrane protein receptor and a therapeutic antibody. As a simple test case, we selected the bactericidal antibody mAB1, which binds to the BamA subunit of the outer membrane-embedded BAM complex ^42^. We purified and reconstituted the 203 kDa BAM complex (comprised of 5 protein subunits, BamA-E) as described above, and analysed successive fractions by MP (Figure 6A and B). Each fraction appeared to contain two overlapping sub-populations: one at 270 kDa and a second less abundant species at 226 kDa (Figure 6B). The mass difference between the two species is ∼40 kDa, which corresponds to the molecular weight of the BamB subunit. Previous work has shown that BamB is prone to dissociating from the full complex after exposure to detergent ^43^. Thus, the 270 kDa population represents the BamABCDE complex reconstituted in peptidiscs, while the 226 kDa population represents a dissociated sub-complex containing BamACDE. Analysis of the antibody mAB1 alone by MP reveals a single species at 156 kDa (Figure 6C).

**Figure 6:**
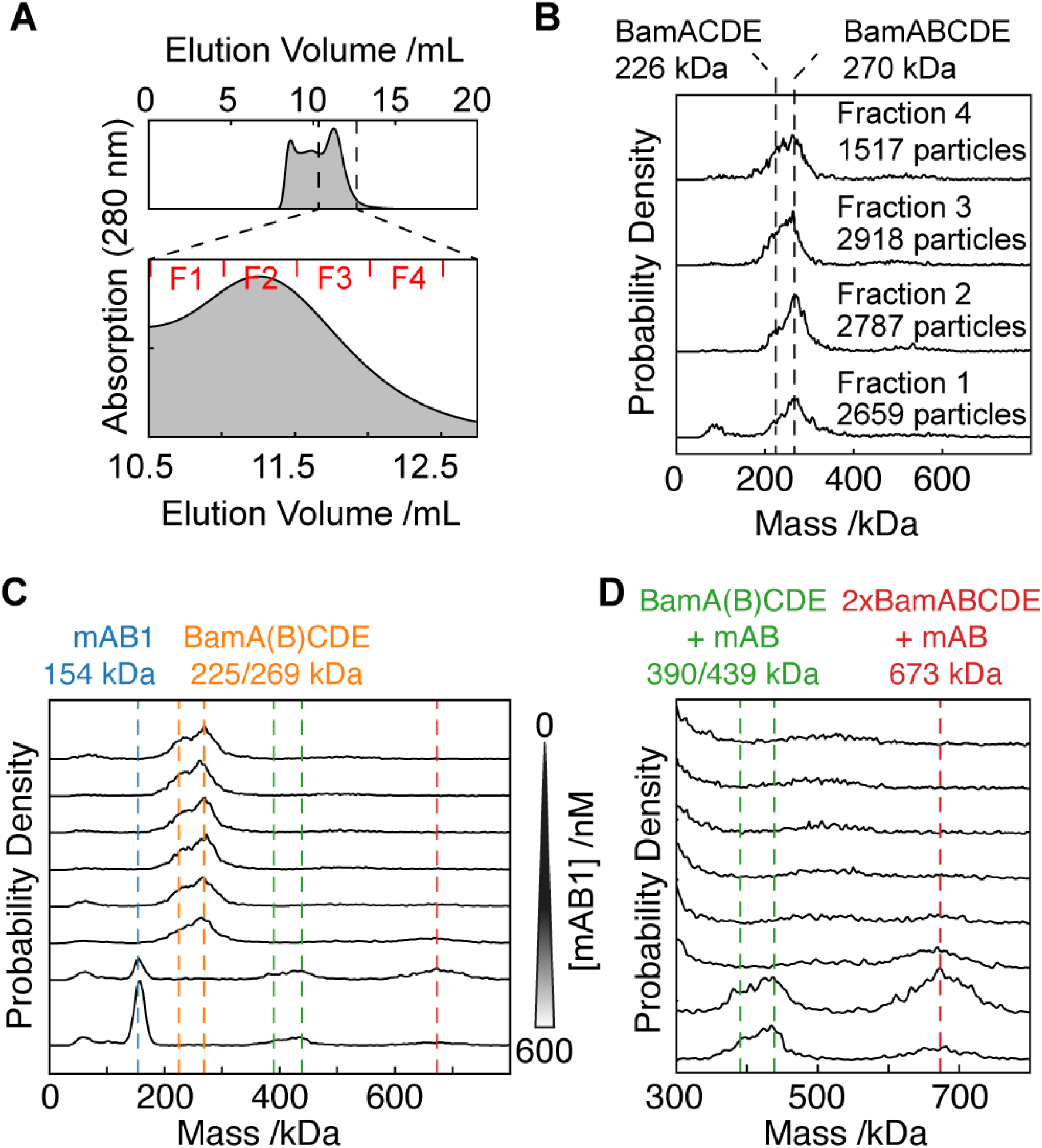
Interaction of the bactericidal antibody mAB1 with the BAM (A and B) Purification and MP analysis of the BAM complex. **(C)** Incubation of the BAM complex (yellow highlight) with increasing concentrations of mAB1 (blue highlight). mAB1 interacts with BAM to form multiple higher-molecular weight species (green and red highlights). **(D)** The lower MW region (below 300 kDa) was cropped from panel **C** to highlight changes in the higher MW range (300 - 800 kDa).

We next performed a titration to visualize the binding of mAB1 with BAM. The concentration of BAM was fixed and the concentration of mAB1 increased from 0 - 600 nM and analysed after 30 min of incubation (Figure 6D). At lower concentrations (0-75 nM) of mAB1, no interaction was evident. At the higher concentrations, however (150-600 nM), we observed two new species at ∼420 and ∼670 kDa. Based on these measured molecular weights, we conclude that the 420 kDa species represents mAB1 binding to one BAM complex, and the 670 kDa species represents mAB1 binding to two BAM complexes (Figure 6D).

## CONCLUSION

In this study, we used Mass Photometry (MP) to characterize bacterial integral membrane proteins reconstituted in peptidiscs. We found that peptidiscs streamlined reconstitution of integral membrane proteins into water soluble particles, requiring only minimal optimization ^9, 12, 22^. Application of our workflow to a series of well-characterized bacterial membrane proteins revealed highly homogenous peptidisc particles (Figure 2, Figure 3A-F). When applied to the tetrameric aquaporin AqpZ and the outer membrane-embedded BAM complex (Figure 3G-I, Figure 6A), we could reveal unexpected sample heterogeneity which was not evident during purification. While it is in principle possible to detect and characterize this heterogeneity using SEC-MALS and negative stain electron microscopy, these techniques are laborious, time-consuming, and require far more protein sample compared to MP ^20, 36^. Furthermore, the resolution obtained here using MP appears much greater than that achievable using either SEC-MALS or electron microscopy.

Beyond screening sample homogeneity, we show, using the ABC transporter BtuCD and the Sec translocon, that MP can be used to quantify high-affinity interactions between membrane protein receptors and their soluble protein ligands (Figure 4, Figure 5). We further used MP to observe binding between the outer membrane-embedded BAM complex and a monoclonal antibody, mAB1, which is bactericidal to *E. coli* under laboratory conditions ^42^. We incubated peptidisc-reconstituted BAM complex with increasing amounts of mAB1 and analysed the mixtures by MP (Figure 6D). At high concentrations of mAB1, we observed formation of two distinct higher-molecular weight species - likely corresponding to mAB1 binding to BAM in a 1:1 and 1:2 ratio. While further experiments will be needed to quantify the kinetics of these two distinct binding events, our results show that MP is a powerful method for characterizing binding of therapeutic antibodies onto membrane protein targets.

Compared with other techniques for quantifying receptor-ligand interactions such as Biolayer Interferometry (BLI), Surface Plasmon Resonance (SPR) and Microscale Thermophoresis (MST), the combination of peptidiscs and MP presented here has several major advantages. Our methodology does not require immobilization of one binding partner onto a chip or surface, as is required in BLI or SPR ^21, 22^, nor is any fluorescent labeling required, which is often needed in MST ^39^. In addition, our method unambiguously reveals the stoichiometry of the observed binding events, which can be very useful for characterizing interactions where previous structural or biochemical data is not available, while also easily distinguishing between native and non-physiological oligomerization. These advantages -coupled with the ease of use, the small amount of valuable biological sample required, and the fact that small molecules which may influence the binding reaction can easily be titrated in - make our approach very useful for *in vitro* screening of antibody-antigen interactions in the rapidly growing field of antibody discovery against membrane protein antigens ^42, 44-47^.

## ACKNOWLEDGEMENTS

We thank Dr. Tarick J. El-Baba and Dr. Jani R. Bolla for helpful discussions during this work. We thank Dr. Steven Rutherford (Genentech) for providing the anti-BamA antibody mAB1. PK is supported by a European Research Council (ERC) Consolidator Grant PHOTOMASS 819593, and an Engineering and Physical Research Council (EPSRC) Leadership Fellowship EP/T03419X/1 (P.K.). Research in the Robinson group is supported By a Wellcome Trust Grant number 221795/Z/20/Z. JWY is supported by a postdoctoral fellowship from the Canadian Institutes of Health Research (MFE-181888). EP is supported by the Deutsche Forschungsgemeinschaft (DFG, German Research Foundation) – project number 455633413.

## CONFLICTS OF INTEREST

P.K. is an academic founder, shareholder, and director to Refeyn Ltd. All other authors declare no conflict of interest.

## SUPPORTING INFORMATION

